# A novel OsIAA3-OsARF16*-OsBUL1* module regulates grain size in rice

**DOI:** 10.1101/2024.09.19.613952

**Authors:** Fengjun Xian, Shuya Liu, Jishuai Huang, Bin Xie, Lin Zhu, Qiannan Zhang, Chen Lv, Yimeng Xu, Xinrong Zhang, Jun Hu

## Abstract

Auxin plays an important role in almost every aspect of plant growth and development. However, the molecular mechanism underlying the control of grain size via auxin signaling pathways is still obscure. Here, we reported that the rice *Aux/IAA* gene *OsIAA3* positively regulates grain size by promoting the cell expansion and proliferation of spikelet hulls. OsIAA3 interacted with 11 OsARFs, among which the interaction with OsARF16 was the strongest. Knockout of *osarf16* led to smaller grain with decreased grain length, grain width, grain thickness and 1,000-grain weight. Meanwhile, transgenic plants overexpressing *OsARF16* produced apparently bigger grain with increased grain length and 1,000-grain weight. Additionally, *OsBUL1*, which positive regulates grain size by promoting cell expansion is a direct target gene of *OsARF16*. Results demonstrated that the interaction between OsIAA3 and OsARF16 repressed the transcriptional activation of OsARF16 on *OsBUL1*. Taken together, our study revealed a novel OsIAA3-OsARF16*-OsBUL1* module which regulates grain size, enriching the molecular mechanism of auxin signaling pathway involved in regulating grain size.

## Introduction

Rice (*Oryza sativa* L.) is a vital staple food for over half the world’s population (Muthayya et al., 2014). Grain yield in rice is determined by three component traits: number of panicles, number of grains per panicle, and grain weight. Grain weight is determined by the grain filling and grain size; with the latter being restricted by the spikelet hull that consists of a palea and a lemma (Li and Li, 2016). A number of genes and quantitative trait loci (QTLs) that associated with grain size have been identified in rice over the past several decades. *GS2*, a positive regulator of grain size and weight, encodes the transcription factor OsGRF4 which regulated by OsmiR396 (Duan et al., 2015). *GS3*, a major QTL for grain length and weight and minor QTL for grain width and thickness, encodes a putative transmembrane protein to negatively regulate grain size (Fan et al., 2006). *GW2*, a QTL for grain width and weight, encodes a functional E3 ubiquitin ligase that ubiquitinates WG1 to negatively regulate grain size (Song et al., 2007; Hao et al., 2021). In rice, OsMKKK10, OsMKK4, and OsMAPK6 act as a cascade that OsMKK4 phosphorylated by OsMKKK10 phosphorylates OsMAPK6 to regulate grain size (Xu et al., 2018). *BG1*, a primary auxin response gene, is involved in the transport of auxin, thereby positively regulating grain size (Liu et al., 2015b). Rice atypical HLH protein *Oryza sativa BRASSINOSTEROID UPREGULATED 1-LIKE* (*OsBUL1*) was reported to as a positive regulator for grain size and leaf angle (Jang et al., 2017). Taken together, grain size is simultaneously controlled by various signaling pathways, including ubiquitin-proteasome pathway, G-protein signaling, mitogen-activated protein kinase (MAPK) signaling, transcriptional regulators and phytohormone signaling (Li et al., 2019). Here, we focused on the regulation of grain size that involved in auxin signaling pathway. The pathway is achieved through the interaction between auxin/indole-3-acetic acid (Aux/IAA) and auxin response transcription factor (ARF) proteins to orchestrate the expression of auxin-dependent gene (Roosjen et al., 2018).

*Aux/IAA* genes, quickly induced by auxin, are classified as early auxin response genes (Walker and Key, 1982; Theologis et al., 1985). These genes encode short-lived nuclear proteins with approximate molecular weights of 20 to 35 kDa, most of which contain four highly conserved domains, designated as Domain I, Domain II, Domain III, and Domain IV (Hagen and Guilfoyle, 2002). Domain I with a LxLxL motif interacts with TOPLESS (TPL), which is important for the function of repression (Tiwari et al., 2004; Szemenyei et al., 2008). Domain II that contains conserved amino acids (GWPPV), is required for the rapid degradation of Aux/IAA proteins (Ramos et al., 2001). Domain III and Domain IV have a similar amino acid sequence to the C-terminal domain of ARF proteins, which mediate homo- and hetero-dimerization among Aux/IAA and ARF proteins (Guilfoyle, 2015).

Auxin response factor (ARF), a DNA binding protein, functions by binding to auxin response elements (AuxREs) in the promoters of auxin response genes, activating or repressing the target genes (Guilfoyle and Hagen, 2007; Guilfoyle, 2015). Most ARF proteins contain an N-terminal B3-type DNA binding domain (DBD), a middle region (MR) and a carboxy-terminal dimerization domain (CTD) (Ulmasov et al., 1999). The DBDs of ARF proteins preferentially bind to AuxREs in the promoters of auxin response genes to regulate their expression (Lieberman-Lazarovich et al., 2019). Interestingly, MRs function as activation domains (ADs) that enriched in glutamine or repression domains (RDs) that lack a glutamine-rich region, which located just CTDs to the DBDs, are not highly conserved in amino acid sequences (Ulmasov et al., 1999). CTDs form homo- and hetero-dimerization among ARF and Aux/IAA proteins, which are not required for binding to AuxREs (Guilfoyle and Hagen, 2001). ARF DBDs alone are sufficient to bind AuxREs and MRs alone are enough for their transcriptional activity, and CTDs along with MRs are required for auxin responses (Tiwari et al., 2003).

In the past decades, many researches have been conducted on the regulation of seed or grain size by Aux/IAA, ARF and Aux/IAA-ARF. In *Arabidopsis*, ARF2/MNT negatively regulates seed size by inhabiting cell division and organ growth (Schruff et al., 2006). In addition, *ARF2* is also involved in ABA-mediated signaling pathway to link the modulation of seed size and drought tolerance (Meng et al., 2015). In *Jatropha curcas*, JcARF19 interacts with JcIAA9 to regulate the seed traits (Ye et al., 2014). In *Brassica napus* L., a 55 amino acids deletion in ARF18 between DBD and MR results in the loss of binding activity, which leads to increased seed weight and silique length (Liu et al., 2015a). In wheat, TaIAA21 interacts with TaARF25 through activating the expression of *TaERF3* to negatively control the grain size (Jia et al., 2021). Loss of function mutations of *TaARF12* lead to significantly increased grain length and 1,000-grain weight (Kong et al., 2023). In rice, the plants of overexpressing *OsARF19* produce thinner grain size (Zhang et al., 2015). OsIAA3 interacts with OsARF25 that positively regulates the grain size by activating the expression level of *OsERF142*/*SMOS1* to negatively regulate grain size, and Gnp4/LAX2, a RING-finger and wd40-associated ubiquitin-like (RAWUL) domain-containing protein, physically interacts with OsIAA3 and consequently interferes with the OsIAA3-OsARF25 interaction (Zhang et al., 2018). *OsAUX3* (the quantitative trait locus *qGL5*) is a negative regulator of grain length by modulating the longitudinal extension of glume cells, and OsARF6 negatively controls grain length by directly binding to the AuxREs of the *OsAUX3* promoter (Qiao et al., 2021). Loss of function mutans of *OsARF4* produces larger rice grain, and OsGSK5/OsSK41 negatively regulates the grain size via phosphorylating OsARF4 (Hu et al., 2018). Recently, a study reported that OsGSK5 interacts with and phosphorylates OsIAA10 to facilitate its interaction with OsTIR1 and subsequent destabilization, however this modification hampers the interaction between OsIAA10 and OsARF4, subsequently, OsARF4 can repress the expression of a positive regulatory gene *OsPHI-1* for grain size (Ma et al., 2023). These researches have greatly enriched our knowledge of seed or grain size regulations by auxin signaling pathways, however, further investigations are still needed to elucidate the molecular network and genetic relationship of seed or grain size controlled by auxin signaling.

Here, we revealed a novel module that OsIAA3-OsARF16*-OsBUL1* for rice grain size regulation. We verified that OsIAA3 interacts with a transcriptional activator OsARF16 to negatively regulate grain size. Then, phenotypic analysis of *OsARF16* transgenic plants revealed that *OsARF16* is a positive regulator for grain size. In our study, we demonstrated that OsARF16 can activate the expression of *OsBUL1* by directly binding to the AuxREs in its promoter. However, the interaction between OsIAA3 and OsARF16 represses this activation. Collectively, these findings define an auxin signaling pathway to regulate grain size promoting the potential application for grain yield.

## Results

### *OsIAA3* negatively regulates grain size

To characterize the biological function of *OsIAA3* in rice, we generated knockout plants of *OsIAA3* using the CRISPR/Cas9 method in the *Oryza sativa* L. *japonica* cultivar Nipponbare. For the *osiaa3-1* mutant, a cytosine (C) located at 438 bp of the *OsIAA3* open reading frame was deleted; for the *osiaa3-2* mutant, an adenine (A) was inserted after 438 bp of the *OsIAA3* open reading frame (Fig. 1A). All of these changes led to early termination of OsIAA3 protein, in which Domain III and Domain IV were lost (Supplementary Fig. S1 and S2A). Meanwhile, we also generated transgenic plants overexpressing *OsIAA3*, *OsIAA3-OE*, driven by the ubiquitin promoter in wild-type background. The qRT-PCR assay validated that the expression levels of *OsIAA3* were significantly higher in *OsIAA3-OE* lines than those in wild-type (Fig. 1B). The examination of these transgenic plants and wild-type showed that both the *osiaa3* mutants produced bigger grain size with increased grain length, grain width and 1,000-grain weight, but no obvious difference of grain thickness were observed when compared to wild-type (Fig. 1, C-G). However, the *OsIAA3-OE* lines, exhibited apparently smaller grain size with decreased grain width, grain thickness and 1,000-grain weight, but no obvious difference of grain length were observed when compared to wild-type (Fig. 1, C-G). These results indicate that *OsIAA3* is a negative regulator of grain size in rice.

**Figure 1.**
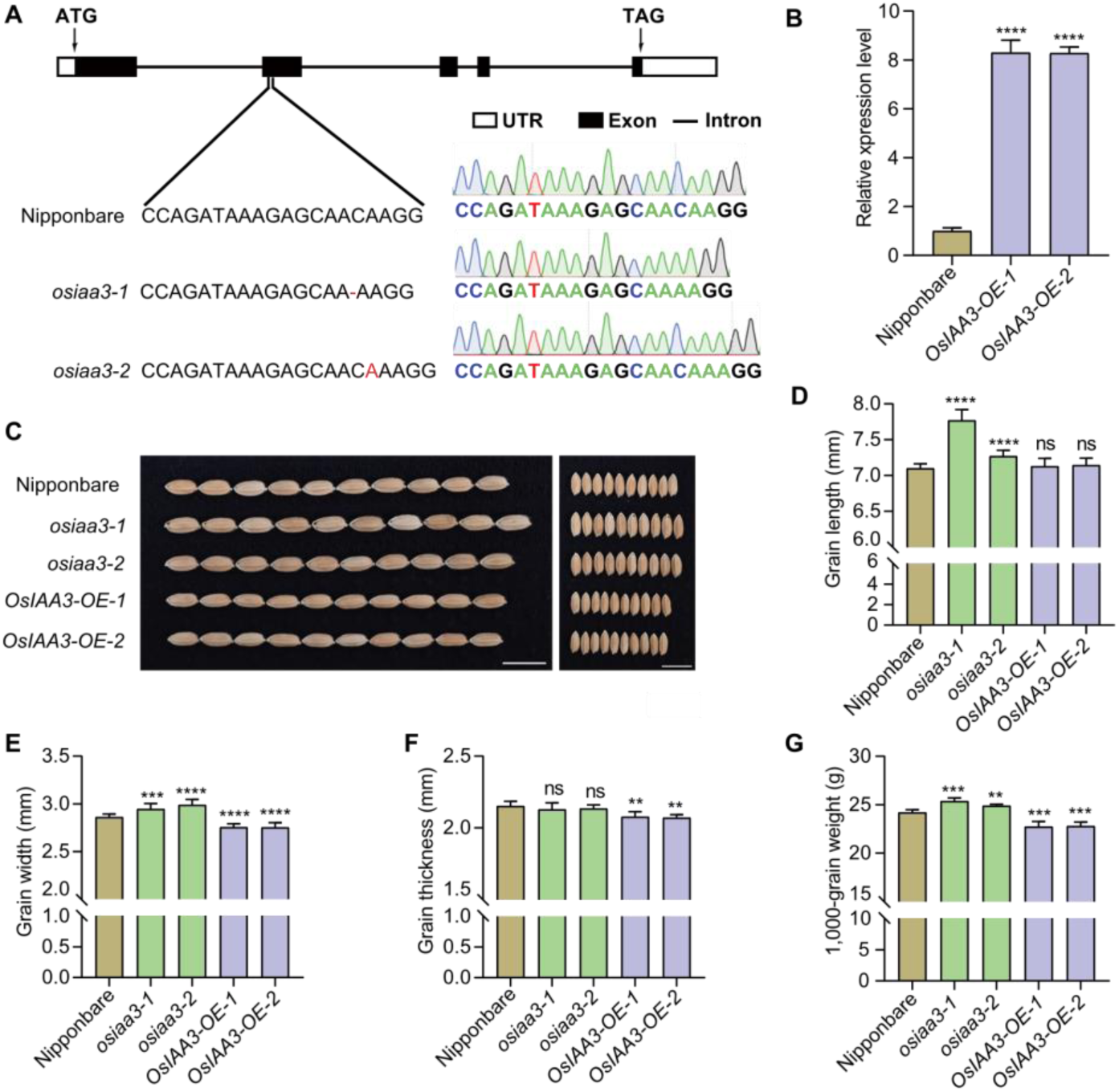
Phenotypic analysis of *OsIAA3* transgenic plants. **(A)** Construction of two *osiaa3* mutants. Black boxes, black lines and white boxes represent exons, introns and UTRs (untranslated regions), respectively. **(B)** Relative expression level of *OsIAA3* in Nipponbare and *OsIAA3-OE* flag leaves at booting stage. Data represent mean values ± SD (*n*=3). The asterisks show significant differences between Nipponbare and *OsIAA3-OE* lines by Student’s *t*-test: ****, *P* < 0.0001. **(C)** Grain morphology of Nipponbare, *osiaa3* and *OsIAA3-OE* plants. Left, grain length; right, grain width. Scale bars, 1cm. **(D-G)** Statistical analysis of grain length, width, thickness and 1,000-grain weight among the lines shown in **(C)**. Data in **(D)** (*n*=10), **(E)** (*n*=10), **(F)** (*n*=5) and **(G)** (*n*=5) are given as means ± SD. Student’s *t*-test was used to generate *P*-values: **, *P* < 0.01; ***, *P* < 0.001; ****, *P* < 0.0001; ns, no significance.

### *OsIAA3* promotes cell expansion and cell proliferation in the spikelet hulls to control grain size

In rice, grain size is mainly restricted by the size of the spikelet hull which is usually associated with cell proliferation and cell expansion (Li and Li, 2016; Li et al., 2019). To get insight into the cytological basis that *OsIAA3* regulates grain size, we observed and compared the outer glume region of mature grains by using scanning electron microscopy (Fig. 2A). We found that the longitudinal epidermal cell length in the *osiaa3* mutant was longer, while which was shorter in *OsIAA3-OE* when compared to wild-type (Fig. 2B). Moreover, the average epidermal cell number per mm^2^ was lower in the *osiaa3* mutant but higher in the *OsIAA3-OE* plants when compared to wild-type (Fig. 2C). Then, we also carried out paraffin section to examine the cell number and area at the cross-section of spikelet hulls (Fig. 2D). Analysis of the hull cross-section in the *osiaa3* mutant revealed significant increases in cell area and cell number within the transverse parenchyma layer (Fig. 2, E-G). In contrast, decreased cell area and cell number were observed in the *OsIAA3-OE* plants (Fig. 2, E-G). Together, these results demonstrated that the bigger grain size of the *osiaa3* mutant is due to the longitudinal elongation and transverse division of the spikelet hulls.

**Figure 2.**
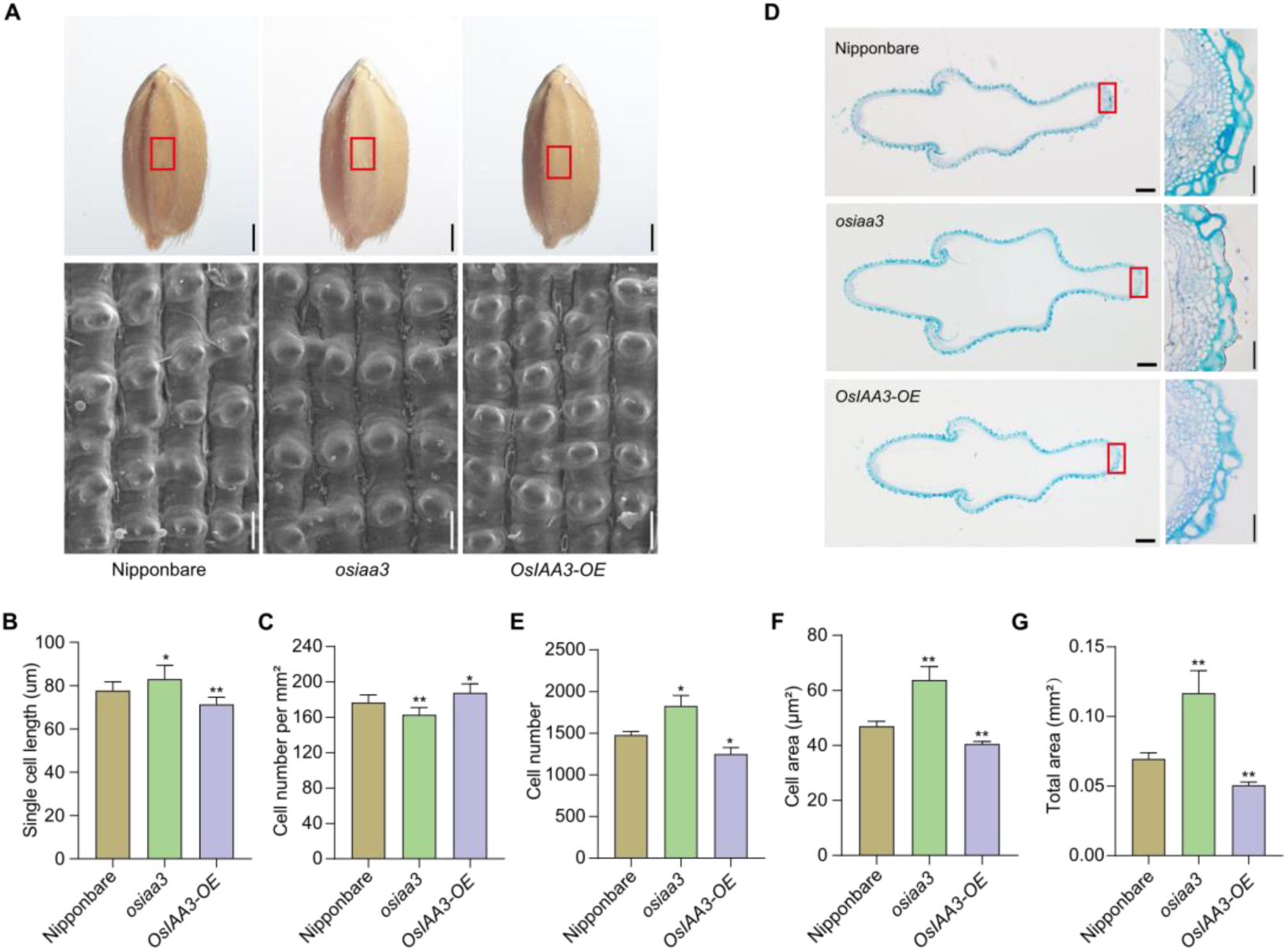
Cytological analysis of glumes in Nipponbare, *osiaa3* and *OsIAA3-OE* plants. **(A)** Scanning electron microscopic analysis of the glume outer surfaces of Nipponbare, *osiaa3* and *OsIAA3-OE* mature grains. Scale bars, 1 mm for whole grains (upper) and 50 μm for outer glume (bottom). **(B-C)** Quantification of single cell length and cell number per mm^2^ in lines shown in **(A)**. Data represent mean values ± SD (*n*=10). The asterisks show significant differences between Nipponbare and transgenic lines by Student’s *t*-test: *, *P* < 0.05; **, *P* < 0.01. **(D)** Cross-sections of spikelet hulls of Nipponbare, *osiaa3* and *OsIAA3-OE* plants. A magnified view of each red boxed cross-section is shown to the right. Scale bars, 200 μm for cross-sections (left) and 30 μm for magnified views (right). **(E-G)** Statistical data of the cell number, cell area and total area in the outer parenchyma layer of the spikelet hulls of Nipponbare, *osiaa3* and *OsIAA3-OE* plants. Data represent mean values ± SD (*n*=3). The asterisks show significant differences between Nipponbare and transgenic lines by Student’s *t*-test: *, *P* < 0.05; **, *P* < 0.01.

### *OsIAA3* is an auxin-responsive gene

*Aux/IAAs*, *SAURs* and *GH3s* are three major classes of early/primary auxin-responsive genes that specifically induced by auxin within minutes (Hagen and Guilfoyle, 2002). To verify whether auxin can induce the expression of *OsIAA3*, 10-day rice seedlings were treated with 1 μM α-Naphthalene acetic acid (NAA). The qRT-PCR results showed that the transcript abundance of *OsIAA3* significantly increased after 4 hours of NAA treatment, and reached the highest level at 8 hours (Fig. 3A). To verify whether auxin induces the degradation of OsIAA3, we expressed the recombinant protein His-OsIAA3 in *E. coli* cells and purified under native conditions. Mixture of the purified protein and total rice protein was treated with 0, 1, 10 and 100 μM concentrations of NAA for 3 h at 30℃. Western blot analysis suggests that the degradation of OsIAA3 by NAA was dose-dependent (Fig. 3B). Moreover, *In vivo* protein degradation assay in rice protoplasts was also carried out. Westen blot results presented that NAA promoted the degradation of OsIAA3, while which was repressed in the presence of MG132 (a specific inhibitor of the 26S proteasome) (Fig. 3C). Together, we concluded that *OsIAA3* is an auxin-responsive gene and is clearly involved in the auxin signaling pathway.

**Figure 3.**
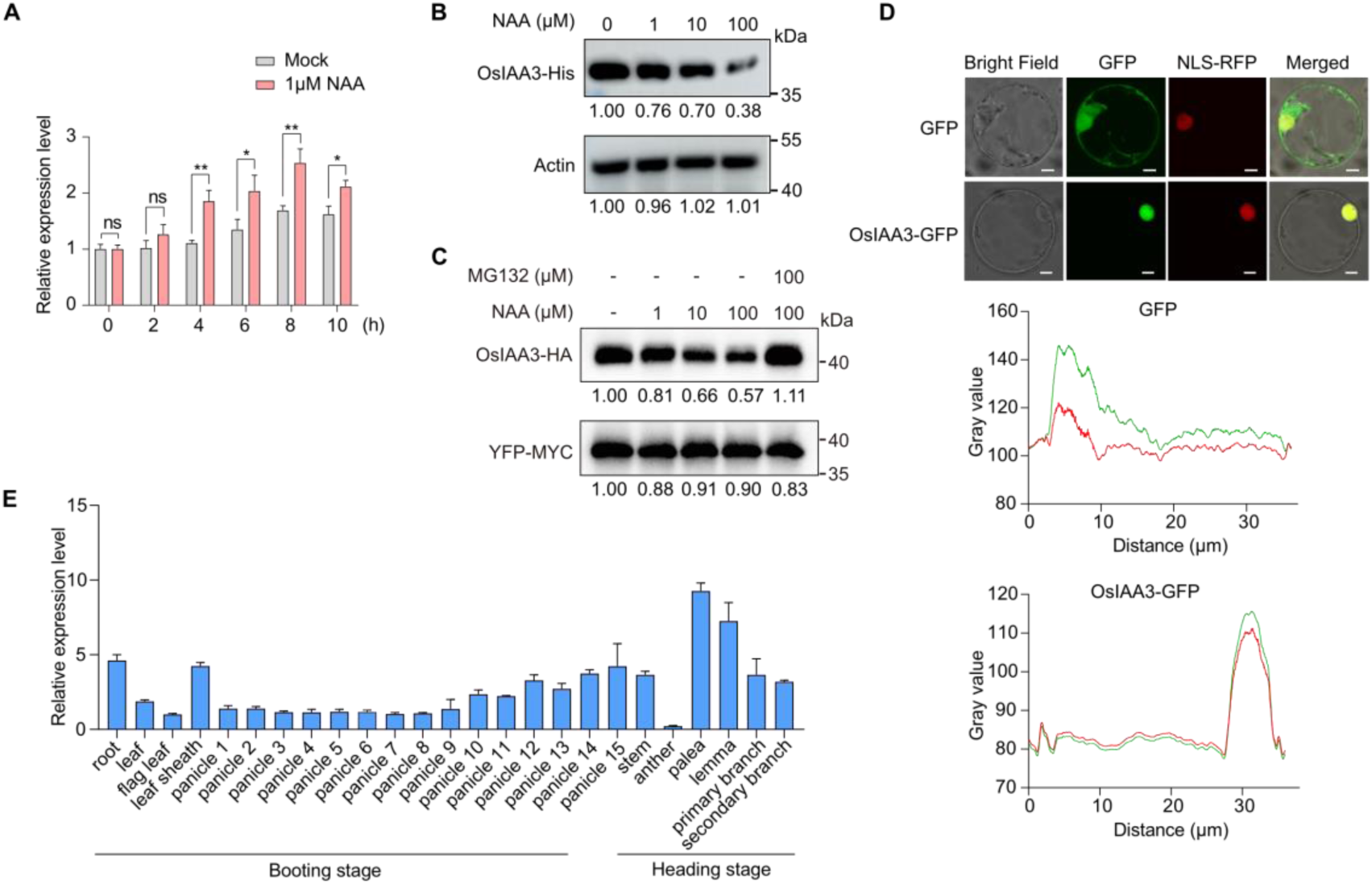
Auxin response, subcellular localization and expression pattern of *OsIAA3*. **(A)** Relative expression level of *OsIAA3* in 10-day rice seedlings treated with 1 μΜ NAA. Data represent mean values ± SD (*n*=3). Student’s *t*-test was used to generate *P*-values: *, *P* < 0.05; **, *P* < 0.01; ns, no significance. **(B)** Cell-free degradation assay of OsIAA3-His protein. The mixture of purified OsIAA3-His protein and rice total protein was treated with different concentrations of NAA at 30℃ for 3 h. Actin protein served as loading control. ImageJ was used to analyze the gray value of each band. **(C)** *In vivo* protein degradation of OsIAA3-HA in rice protoplasts. YFP-MYC recombination protein was used as control. ImageJ was used to analyze the gray value of each band. **(D)** Subcellular localization of OsIAA3-GFP fusion protein in rice protoplasts. D53-mCherry was used as a nuclear marker (upper). Scale bars, 5 μm. The pixel intensities for the different color channels along transects was conducted by ImageJ (bottom). **(E)** Relative expression level of *OsIAA3* in various tissues and stages of Nipponbare. Three independent biological replicates were used in this experiment. Panicle 1 represent young panicle about 0-1 cm, panicle 2 represent young panicle about 1-2 cm, panicle 3 represent young panicle about 2-3 cm and so on.

### OsIAA3 is localized in nucleus and highly expressed in palea and lemma

To investigate the subcellular localization of the OsIAA3, the CDS of *OsIAA3* was fused with GFP for transient expression in rice protoplast. Then OsIAA3-GFP construct was co-transformed with a nuclear localization marker (NLS-RFP) simultaneously. We observed that the fluorescence of the OsIAA3-GFP was co-localized with NLS-RFP in the nucleus, indicating the nuclear localization of OsIAA3 (Fig. 3D). Bioinformatics analysis has predicted that both Domains II and Domain IV contain putative nuclear localization signals (NLSs) in rice (Jain et al., 2006). To determine which domain contributes to the subcellular localization of OsIAA3, we constructed a series of GFP fusion vectors according to the distribution of the domains of OsIAA3 (Supplementary Fig. S2A). We observed that, Domain II was sufficient to drive the protein localization, rather than Domain IV (Supplementary Fig. S2B). Aux/IAA proteins can form homodimers via conserved Domain III and Domain IV. To confirm whether OsIAA3 can form homodimers, we performed yeast two-hybrid (Y2H) assay using the intact and truncated protein, respectively. These results demonstrated that intact OsIAA3 protein can form homodimers, and the presence of Domain III is important for this homodimerization process (Supplementary Fig. S2C).

To examine the expression pattern of *OsIAA3* in wild-type, we performed qRT-PCR assay with various tissues, including root, leaf, flag leaf, leaf sheath, panicles at the booting stage and so on. Results showed that *OsIAA3* was expressed ubiquitously, except lower in anther (Fig. 3E), suggesting its various functions. It is worth mentioning that *OsIAA3* was highly expressed in palea and lemma, suggesting that *OsIAA3* might play an important role in spikelet hull development, consistent with our previous study. These results imply that *OsIAA3* plays an important role in regulating grain size.

### OsIAA3 interacts with 11 OsARF proteins

Aux/IAA proteins regulate the expression of auxin-responsive gene by repressing the activity of ARF proteins (Guilfoyle, 2015). The rice genome encodes 25 OsARF proteins, of which 9 are transcriptional activators and 16 are transcriptional repressors (Shen et al., 2010). In previous report, OsARF activators, but not the repressors, interact with Aux/IAA proteins in Y2H assay (Shen et al., 2010). To investigate the interaction network between OsIAA3 and OsARFs, we cloned 22 *OsARF* genes, and tested the interactions by Y2H assay point to point. We observed that several OsARFs, including 1, 5, 6, 7, 11, 12, 16, 17, 19, 21 and 25, interacted with OsIAA3 (Fig. 4A and Supplementary Fig. S3). However, the interaction intensity was strongest between OsIAA3 and OsARF16 than others (Fig. 4A). To further validate these interactions in plants, we conducted a firefly luciferase complementation imaging (LCI) assay. Luminescence signals were detected in cLUC-OsARF1, cLUC-OsARF5, cLUC-OsARF6, cLUC-OsARF7, cLUC-OsARF11, cLUC-OsARF12, cLUC-OsARF16, cLUC-OsARF17, cLUC-OsARF19, cLUC-OsARF21, cLUC-OsARF25 and nLUC-OsIAA3 co-expressing regions (Fig. 4B). The signals were not observed between other cLUC-OsARFs and nLUC-OsIAA3 co-expressing region, as well as negative control (Supplementary Fig. S4). Regarding to the strongest interaction between OsARF16 and OsIAA3, we also confirmed the interaction using bimolecular fluorescence complementation (BiFC) assay. The fluorescence signals from SPYNE-OsARF16 and SPYCE-OsIAA3 were clearly observed, while those from the negative control were not (Supplementary Fig. S5). Taken together, OsIAA3 interacts with 11 OsARFs, 2 repressors (OsARF1 and OsARF7) and 9 activators, among which the interaction between OsIAA3 and OsARF16 is highly solid. Therefore, we firstly focused on *OsARF16* for further study.

**Figure 4.**
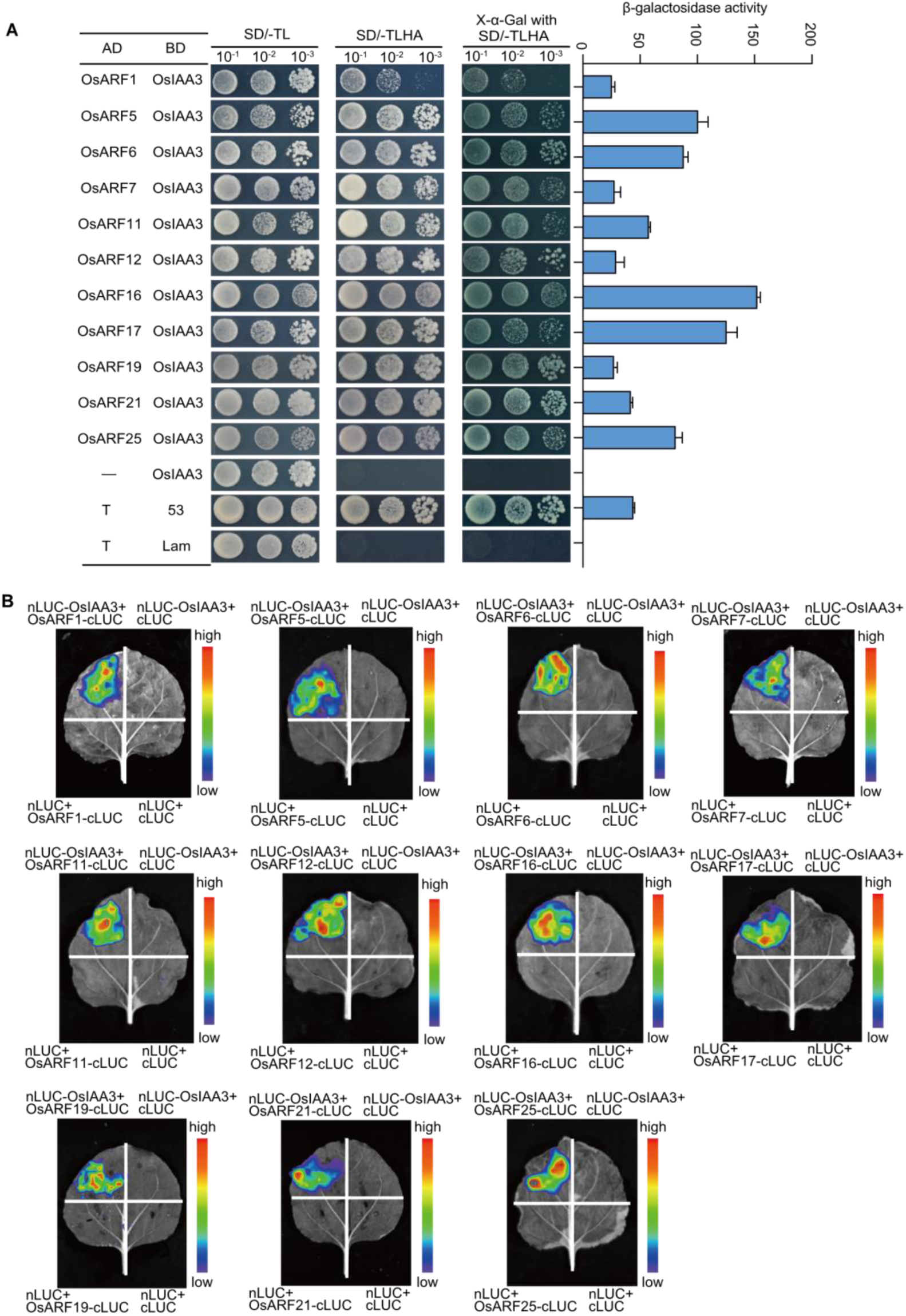
Interaction analysis of OsIAA3 and 11 OsARFs. **(A)** Y2H assay to detect the interactions of OsIAA3 and 11 OsARFs (left). β-galactosidase activity was measured to evaluate the intensity of interactions (right) (*n*=3). The interaction of pGBKT7-53 with pGADT7-T was used as positive control, and the interaction of pGBKT7-Lam with pGADT7-T was used as negative control. **(B)** Firefly LCI assay to detect the interactions of OsIAA3 and 11 OsARFs in *Nicotiana benthamiana*.

### OsARF16, a transcriptional activator, is highly expressed in palea, lemma and young panicles

To investigate the transcriptional activation of OsARF16 in yeast, we fused the intact and truncated *OsARF16* fragments in the pGBKT7 vector, respectively. The recombinant vectors were separately co-transformed with pGADT7 into the yeast strain AH109. We observed that intact OsARF16 had strong transactivation activity, while the truncated OsARF16 had no transactivation activity (Fig. 5, A and B). To test the transcriptional activity in plants, we next fused *OsARF16* fragment in GAL4DBD vector to generate the construct GAL4DBD-OsARF16. The *luciferase* contained five copies of binding sites for GAL4 was used as a reporter, and the *Renilla luciferase* was used as an internal control (Fig. 5C). Compared with the GAL4DBD empty vector, the LUC/REN ratio was significantly increased in protoplasts expressing GAL4DBD-OsARF16 plasmid (Fig. 5D). These results confirmed that OsARF16 acts as a transcriptional activator in rice.

**Figure 5.**
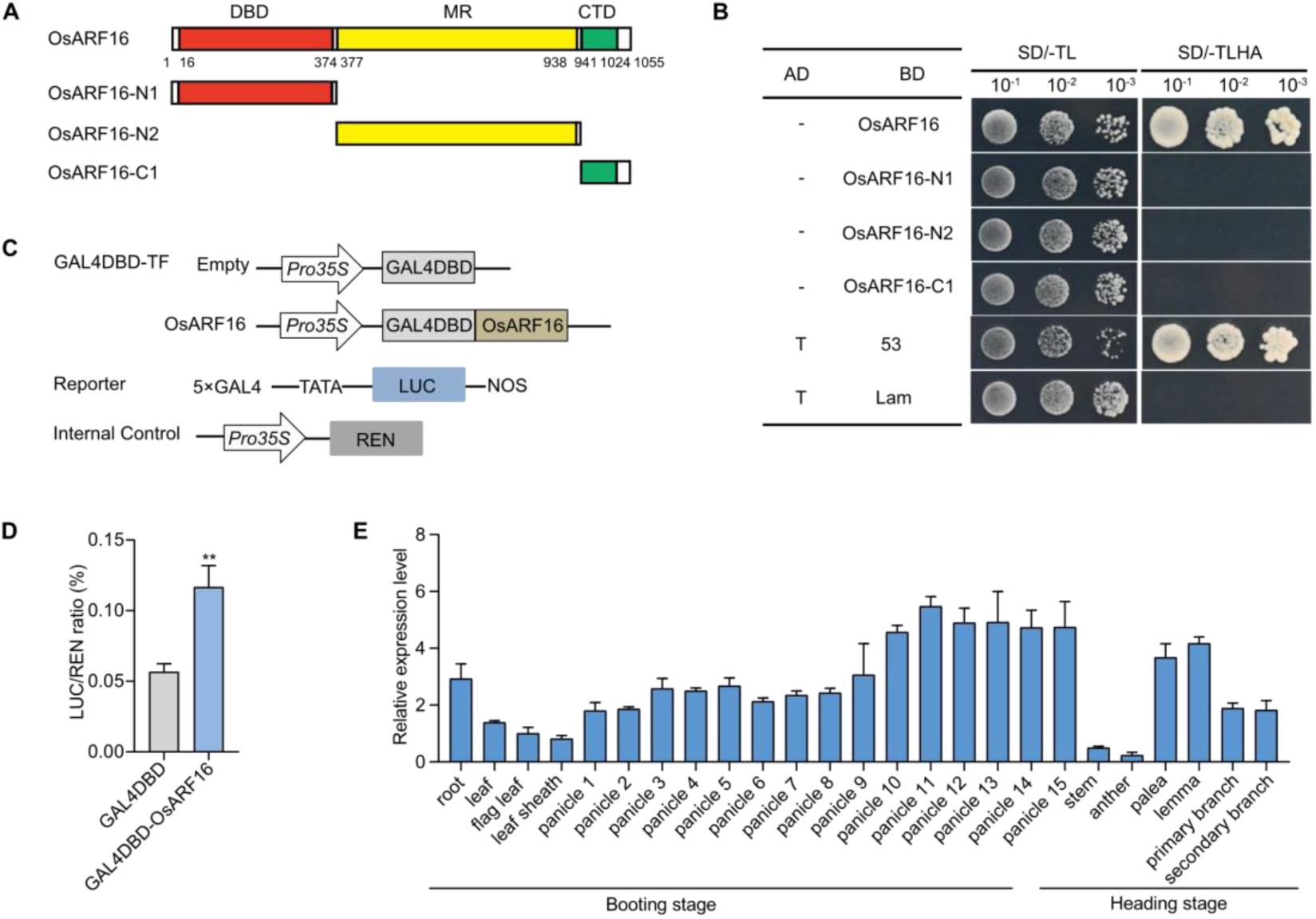
Transcriptional activity and expression pattern analysis of *OsARF16*. **(A)** The modular structure of OsARF16. DBD, DNA-binding domain; MR, middle region; CTD, C-terminal dimerization domain. **(B)** Transcriptional activity analysis of OsARF16 in yeast. pGBKT7-53 with pGADT7-T was used as positive control, and pGBKT7-Lam with pGADT7-T was used as negative control. **(C)** Schematic diagrams of GAL4DBD-TF, reporter and internal control used for transcriptional activation analysis. **(D)** The transcriptional activity assay of OsARF16 in rice protoplasts. Data represent mean values ± SD (*n*=3). Student’s *t*-test was used to generate *P*-values: **, *P* < 0.01. **(E)** Relative expression level of *OsARF16* in various tissues and stages of Nipponbare. Three independent biological replicates were used in this experiment. Panicle 1 represent young panicle about 0-1 cm, panicle 2 represent young panicle about 1-2 cm, panicle 3 represent young panicle about 2-3 cm and so on.

Moreover, expression pattern analysis indicated that *OsARF16* constitutively expressed in all tissues, including the roots, stems, leaves and panicles (Fig. 5E). Interestingly, *OsARF16* was highly expressed in palea, lemma and young panicles, which was similar to the expression pattern of *OsIAA3*. Therefore, it strengthened our speculation that OsIAA3 regulates the grain size through interacting with OsARF16.

### *OsARF16* positively regulates grain size

To further explore the biological function of *OsARF16*, the CRISPR/Cas9 system was employed to create knockout plants in the Nipponbare background. The *osarf16* mutant, a thymine (T) was inserted after 117 bp of the *OsARF16* open reading frame, which led to an early termination of OsARF16 protein (Fig. 6A and Supplementary Fig. S6). Besides, we also performed overexpression plants of *OsARF16, OsARF16-OE*, driven by the ubiquitin promoter in the wild-type background. The expression levels of *OsARF16* were significantly higher in the *OsARF16-OE* plants than in wild-type (Fig. 6B). The *osarf16* mutant produced obviously smaller grain size with decreased grain length, grain width, grain thickness and 1,000-grain weight when compared to wild-type (Fig. 6, D-G). However, the *OsARF16-OE* lines, exhibited apparently bigger grain size with increased grain length and 1,000-grain weight when compared to wild-type (Fig. 6, D-G). These results suggest that *OsARF16* is a positive regulator of grain size in rice.

**Figure 6.**
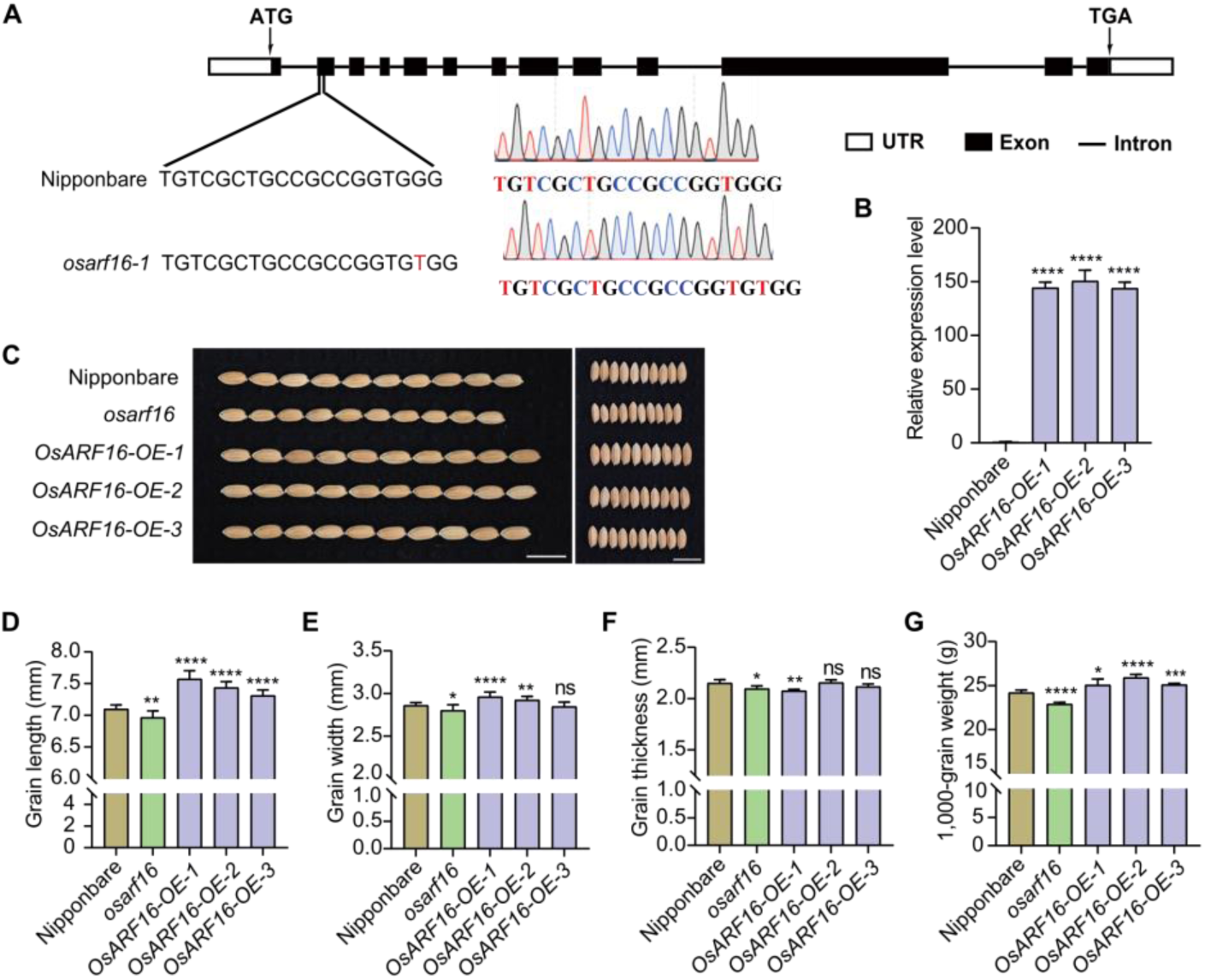
Phenotypic analysis of *OsARF16* transgenic plants. **(A)** Identification of *osarf16*. Black boxes, black lines and white boxes represent exons, introns and UTRs (untranslated regions), respectively. **(B)** Relative expression of *OsARF16* in Nipponbare and *OsARF16-OE* leaves at booting stage. Data represent mean values ± SD (*n*=3). The asterisks show significant differences between Nipponbare and *OsARF16-OE* lines by Student’s *t*-test: ****, *P* < 0.0001. **(C)** Grain morphology of Nipponbare, *osarf16* and *OsARF16-OE* plants. Left, grain length; right, grain width. Scale bars, 1cm. **(D-G)** Statistical analysis of grain length, width, thickness and 1,000-grain weight among the lines shown in **(C)**. Data in **(D)** (*n*=10), **(E)** (*n*=10), **(F)** (*n*=5) and **(G)** (*n*=5) are given as means ± SD. Student’s *t*-test was used to generate *P*-values: *, *P* < 0.05; **, *P* < 0.01; ***, *P* < 0.001; ****, *P* < 0.0001; ns, no significance.

Next, we observed and compared the cell length and cell number per mm^2^ of spikelet hulls by using scanning electron microscopy (Supplementary Fig. S7A). We found that the epidermal single cell length of the spikelet hull in the *osarf16* mutant was shorter, while which was longer in the *OsARF16-OE* lines when compared to wild-type (Supplementary Fig. S7B). Moreover, the average epidermal cell number per mm^2^ was higher in the *osarf16* mutant but lower in the *OsARF16-OE* lines when compared to wild-type (Supplementary Fig. S7C). Paraffin section assay also revealed a significantly decreased cell area and cell number in the *osarf16* mutant, while increased cell area and cell number were observed in the *OsARF16-OE* lines (Supplementary Fig. S7, D-G). Subsequently, we tested the relative expression of several cell cycle-regulated genes in the *OsARF16* transgenic plants. Consistent with these observations, 9 cell cycle-related genes were up-regulated in the *OsARF16-OE* plants, and 7 of which were down-regulated in the *osarf16* mutant (Supplementary Fig. S8). These results suggest *OsARF16* regulates grain size through promoting cell expansion and cell proliferation in the spikelet hulls, which shared a similar cytological basis with *OsIAA3*.

### OsARF16 regulates grain size by activating the expression of *OsBUL1*

To explore the underlying molecular mechanisms by which *OsARF16* regulates grain size, RNA sequencing (RNA-seq) analysis was performed to analyze the transcriptome of both *OsARF16-OE* and wild-type plants. Compared with wild-type, a total of 2166 differentially expressed genes (DEGs) were detected, among which 1794 genes were up-regulated and 372 genes were down-regulated in the *OsARF16-OE* plants (Supplementary Fig. S9 and Supplementary table2). We focused on the differentially expressed up-regulated genes regulating grain size in the *OsARF16-OE* plants. Then, 16 genes were selected as candidate genes, and we detected the expression level of these genes in *OsARF16* transgenic plants by qRT-PCR. The results showed that 5 genes down-regulated in the *osarf16* mutant while up-regulated in the *OsARF16-OE* lines compared to those in wild-type (Fig. 7A). Motif scanning analysis of these 5 candidate genes revealed the presence of one AuxRE (TGTCTC) in the promoter region of *DRW1* and three AuxREs in the promoter regions of *OsBUL1*, *OsOPL1* and *OsSPL13* (Fig. 7B and Supplementary Fig. S10A), implying that they are potential downstream targets of *OsARF16*. To test whether OsARF16 directly regulates the expression of these 4 genes, 2-kb promoter fragments of those genes were fused to the luciferase reporter gene coding sequence (Fig. 7C and Supplementary Fig. S10B), and performed expression experiments in rice protoplasts and *Nicotiana benthamiana*, simultaneously. The LUC/REN ratio was significantly increased when *OsARF16* was co-expressed with *ProOsBUL1* reporter but not with *ProDRW1*, *ProOsOPL1* and *ProOsSPL13* reporters (Fig. 7D and Supplementary Fig. S10C), which was consistent with the observations in *Nicotiana benthamiana* (Fig. 7E and Supplementary Fig. S10D). In addition, electrophoretic mobility shift assay (EMSA) further validated that recombinant GST-OsARF16^114-237^ protein caused shift bands of the FAM-labeled probes (Fig. 7F). The shifted bands were significantly reduced in the presence of non-labeled (cold) probes with a dosage-dependent manner (Fig. 7F). All of these results suggest that OsARF16 regulates the expression of *OsBUL1* by directly binding to its promoters. In addition, qRT-PCR were also carried out to investigate the expression pattern of *OsBUL1*. Results revealed that *OsBUL1* was highly expressed in palea, lemma and young panicles (Supplementary Fig. S11), which was quite similar to the expression pattern of *OsIAA3* and *OsARF16*. These results established OsIAA3 regulates grain size via the OsARF16*-OsBUL1* module.

**Figure 7.**
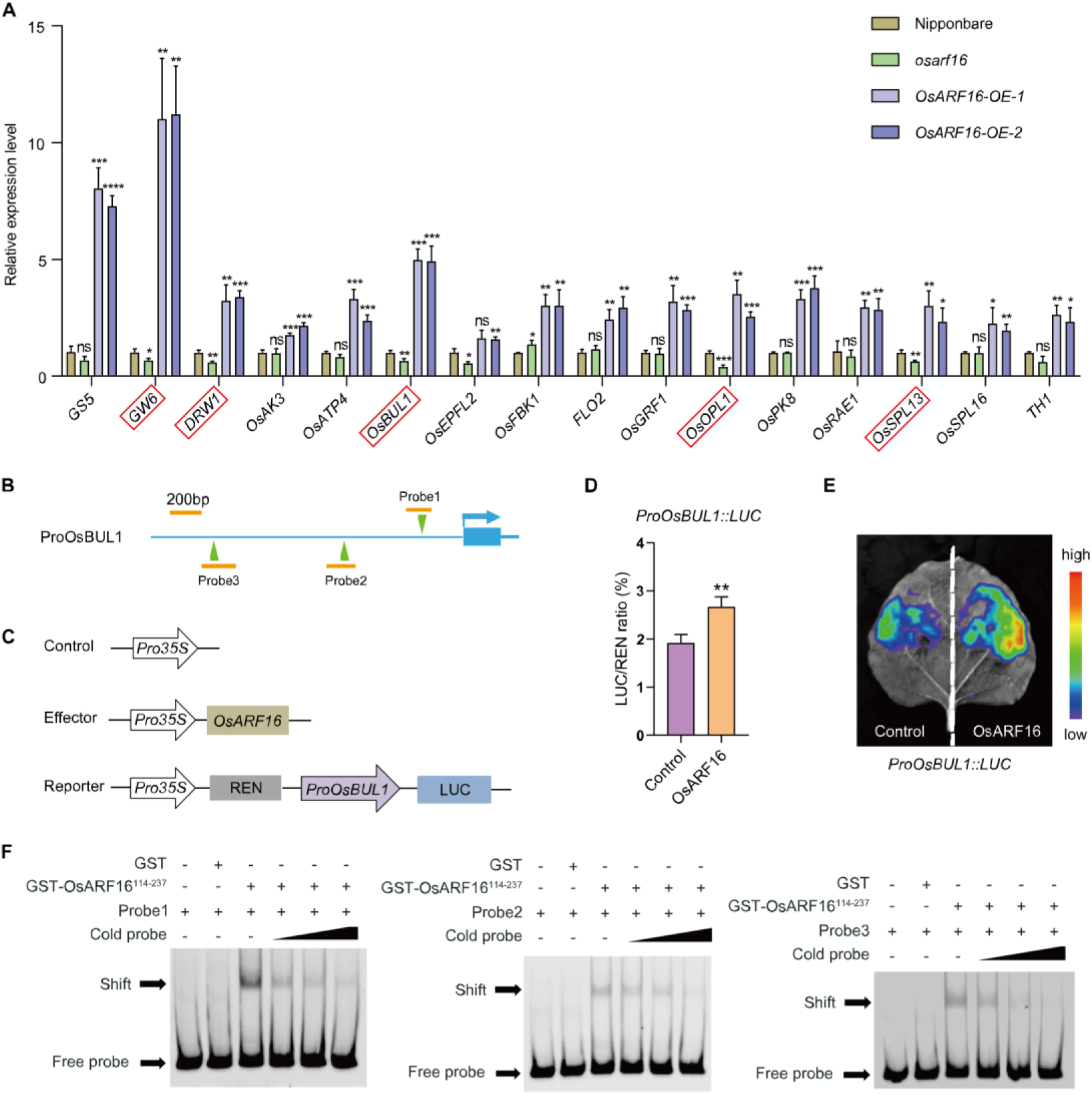
OsARF16 activates the expression level of *OsBUL1*. **(A)** Relative expression levels of grain-size genes in Nipponbare, *osarf16* and *OsARF16-OE* plants. Data represent mean values ± SD (*n*=3). Student’s *t*-test was used to generate *P*-values: *, *P* < 0.05; **, *P* < 0.01; ***, *P* < 0.001; ****, *P* < 0.0001; ns, no significance. **(B)** Schematic diagrams of the AuxREs (TGTCTC) in the promoter region of *OsBUL1*. The green triangles and arrows respectively indicate the position and direction of AuxREs in the promoter region. The orange lines represent the amplified regions for EMSA analysis. **(C)** Schematic diagrams of the control, effector and reporter used for dual-luciferase transcriptional activity assay. **(D-E)** The transient dual-luciferase transcriptional activity assay of OsARF16 in rice protoplasts **(D)** and *Nicotiana benthamiana* **(E)**. Data represent mean values ± SD (*n*=3). Student’s *t*-test was used to generate *P*-values: **, *P* < 0.01. **(F)** EMSA analysis of OsARF16 binding to the AuxREs in the *OsBUL1* promoter. The probes were designed in the *OsBUL1* promoter **(B)**. GST protein was used as negative control. Two-fold, five-fold and ten-fold unlabeled probes were used as cold probes. + and – indicate the presence and absence of the related probes and probes.

### OsIAA3 represses the transcriptional activation of OsARF16

As transcriptional repressors in the auxin signaling pathway, Aux/IAA proteins interact with ARFs to repress the transcriptional activity of ARFs. To determine whether OsIAA3 represses the transcriptional activation of OsARF16, we co-transformed GAL4DBD-OsARF16 with the effector OsIAA3 or empty vector (negative control) in rice protoplasts (Fig. 8A). The LUC/REN ratio of GAL4DBD-OsARF16 was dramatically decreased when co-expressed with OsIAA3, as compared to when it co-expressed with the empty vector (Fig. 8B), suggesting the transactivation of OsARF16 is repressed by OsIAA3. To further investigate whether OsIAA3 inhibits the transcriptional regulation of OsARF16 on *OsBUL1*, we co-transformed the reporter ProOsBUL1::LUC, the effectors OsARF16 with or without OsIAA3 (Fig. 8C). Results showed that the LUC/REN ratio of ProOsBUL1::LUC was significantly decreased when co-expressed with the effector OsARF16 with OsIAA3 (Fig. 8D). Taken together, these results suggest that OsIAA3 represses the transcriptional activation of OsARF16 on *OsBUL1*.

**Figure 8.**
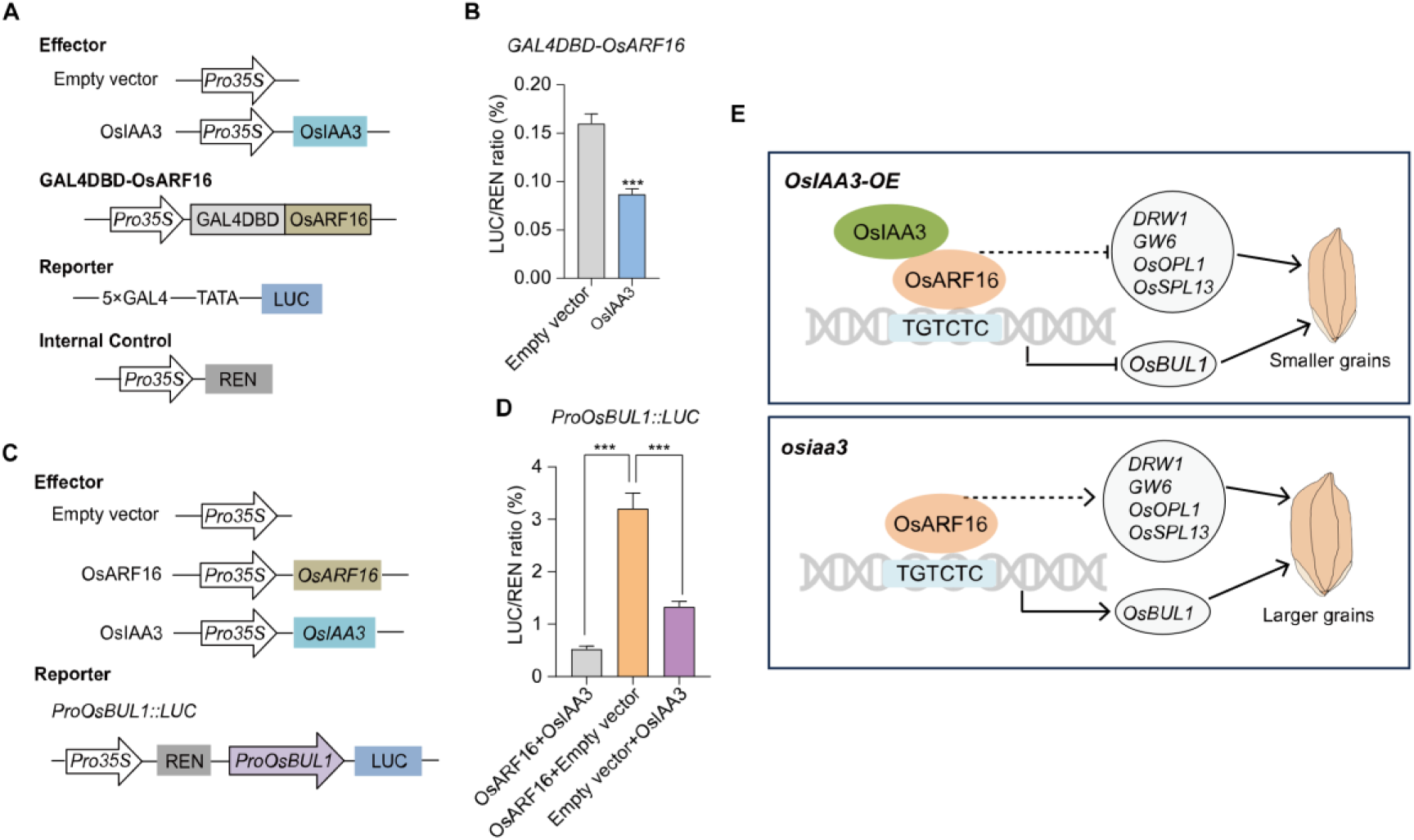
OsIAA3 represses the transcriptional activation of OsARF16. **(A)** Schematic diagrams of effector, GAL4DBD-TF, reporter and internal control used for transcriptional analysis. **(B)** The transcriptional activity assay of OsARF16 in rice protoplasts. Data represent mean values ± SD (*n*=3). Student’s *t*-test was used to generate *P*-values: ***, *P* < 0.001. **(C)** Schematic diagrams of the effector and reporter used for dual-luciferase transcriptional activity assay. **(D)** The transient dual-luciferase transcriptional activity assay of OsARF16 in rice protoplasts. Data represent mean values ± SD (*n*=3). Student’s *t*-test was used to generate *P*-values: ***, *P* < 0.001. **(E)** A proposed model of the OsIAA3-OsARF16*-OsBUL1* module for the regulation of grain size in rice. OsAR16 indirectly promotes the expressions of positive regulatory genes (*DRW1*, *GW6*, *OsOPL1* and *OsSPL13*) associated with grain size and directly activates the expression of *OsBUL1* by binding to the AuxREs (TGTCTC) in the promoter region, ultimately leading to an increase in rice grain size. However, the transcriptional activation of OsARF16 on *OsBUL1* is repressed by the interaction between OsARF16 and OsIAA3.

Given that the axis of OsIAA3*-*OsARF16*-OsBUL1*, to verify whether *OsBUL1* is involved in auxin signaling pathway, 10-day seedlings of the *osarf16* mutant and wild-type were treated with 1 μM NAA. The qRT-PCR showed that the expression level of *OsBUL1* was significantly induced by NAA in wild-type, while not in the *osarf16* mutant (Supplementary Fig. S12). This result suggests auxin can regulate *OsBUL1* through OsIAA3-OsARF16 module. Therefore, an auxin signaling pathway was uncovered, in which the auxin-OsIAA3-OsARF16-*OsBUL1* axis is essential for grain size control by regulating cell expansion and cell proliferation in the spikelet hulls.

## Discussion

### OsIAA3 regulates grain size via interaction with OsARF16

Auxin plays a pivotal role in almost every aspect of plant growth and development, and the Aux/IAA and ARF proteins are key regulators in auxin signaling pathway. In our study, we shown that *OsIAA3* negatively regulates grain size by promoting cell expansion and proliferation in spikelet hulls (Fig. 1-2). Aux/IAA proteins are short-lived nuclear proteins that repress the expression of auxin response genes through interaction with ARF proteins (Tiwari et al., 2004). OsIAA3 interacted with 11 OsARFs, with the strongest interaction observed in association with OsARF16 (Fig. 4 and Supplementary Fig. S5). Certainly, we are uncovering the functions of other *OsARFs* in our lab (data not shown). Moreover, expression pattern analysis showed that *OsIAA3* and *OsARF16* are highly expressed in palea, lemma and young panicles (Fig. 3E and 5E). Subsequently, CRISPR/Cas9 technology was employed to generate the loss of function mutant, *osarf16*, which exhibited smaller grain size. In contrast, plants overexpressing *OsARF16* (*OsARF16-OE*) displayed phenotypes that were converse of those observed in the *osarf16* mutant. These results revealed that *OsIAA3* and *OsARF16* regulate grain size via a common pathway.

### *OsBUL1* is one of the target genes of OsARF16 to regulate grain size

The *drw1* mutant exhibits a severe dwarf phenotype with smaller grain size compared to the wild-type ZH11 (Zhang et al., 2021). Overexpression of *OsOPL1* leads to larger grain size, which could be attributed to the enhanced proliferation of cells (Zhang et al., 2022). *GW6* is reported to positively regulate grain width and weight by promoting cell expansion in the spikelet hulls (Shi et al., 2020). *OsSPL13* (also named *GLW7*) positively regulates grain length and yield by promoting cell expansion in the spikelet hulls (Si et al., 2016). *Oryza sativa* BRASSINOSTEROID UPREGULATED 1-LIKE1 (OsBUL1) is an atypical HLH protein, which positively regulates grain size by promoting cell expansion (Heang and Sassa, 2012; Jang et al., 2017). In our study, RNA-seq analysis revealed a series of grain-size genes up-regulated in the *OsARF16-OE* plants, and qRT-PCR verified that *DRW1*, *OsOPL1*, *GW6*, *OsSPL13*, and *OsBUL1* down-regulated in the *osarf16* mutant (Fig. 7A). *OsARF16* positively regulated grain size by promoting cell expansion and proliferation in the spikelet hulls (Supplementary Fig. S7). Dual-luciferase transcriptional activity assay and EMSA indicated OsARF16 directly binds to the AuxRE of *OsBUL1* promoter and activates the expression of *OsBUL1* (Fig. 7, B-F). Therefore, further investigation is expected to elucidate the regulatory mechanisms by which OsARF16 modulates the expression of other four genes. These results indicated *OsARF16*, acting as a transcriptional activator, regulates grain size by directly regulating *OsBUL1* and indirectly regulating other four grain-size genes.

### Aux/IAA and ARF interaction network

Aux/IAA proteins act as repressors by directly interacting with ARF proteins and recruiting the co-repressors protein TOPLESS (TPL) and TOPLESS RELATED (TPR) through an EAR motif to the chromatin (Szemenyei et al., 2008). At high auxin levels, Aux/IAA proteins are bound by SCF^TIR1^ and degraded through the ubiquitin-proteasome pathway, which relieves the suppression of ARF proteins (Gray et al., 2001; Guilfoyle, 2015; Salehin et al., 2015). The rice genome encodes 25 OsARF proteins and 31 Aux/IAA proteins, and the interaction network between 14 OsARF and 15 OsIAA proteins were investigated previously (Shen et al., 2010). The complicated interaction network implies abundant functions and fine regulations between Aux/IAA and ARF proteins. In this study, OsIAA3 interacted with OsARF1, 5, 6, 7, 11, 12, 16, 17, 19, 21, 25 (Fig. 4). Of which, OsARF1 interacts with OsIAA6 to negatively regulate leaf inclination, and overexpression of *OsARF1* results in the recovered leaf inclination of *osiaa6-D* (a gain-of function mutant) (Xing et al., 2022). OsARF17 interacts with OsIAA12 to regulate inclination, and overexpression of *OsIAA12* or deficiency of *OsARF17* results in enlarged leaf inclination (Chen et al., 2018). OsARF19 interacts with OsIAA13 and directly regulates *LBD1-8* expression, which modulates lateral roots formation (Yamauchi et al., 2019). Here, we revealed a novel OsIAA3-OsARF16 module to regulate *OsBUL1* for grain size. However, the functions between OsIAA3 and other OsARF proteins still require further study. On the basis of these findings, we propose a working model for OsIAA3*-*OsARF16*-OsBUL1* in rice. Regarding to the auxin signaling pathway contributing to grain size, OsIAA3 was fine adjusted by endogenous auxin. OsIAA3, a negative regulator of grain size, interacts with OsARF16 and represses the transcriptional activation of OsARF16 on *OsBUL1* and other unknown genes (Fig. 8E). The increased expression level of *OsBUL1* and several grain-size genes promotes the cell expansion and cell proliferation of spikelet hulls, resulting in larger grains. These findings expand recent knowledge how auxin regulates the grain size in rice and provide an alternative strategy for improving yield in the future.

## Materials and Methods

### Plant materials and phenotypic analysis

Rice *Oryza sativa* L*. japonica* cultivar Nipponbare was used as the transgenic receptor in this study. All the rice plants were grown in paddy fields in either Wuhan (May to October) or Hainan (December to May), China. The plant agronomic traits were collected in Wuhan under proper management. At least 10 plants were harvested to measure the grain length, grain width, grain thickness and 1,000-grain weight. *Nicotiana benthamiana* plants were grown in a greenhouse at 22℃ with 60% relative humidity and a 16 h light/8 h dark photoperiod at Wuhan University.

### Vector construction and plant transformation

The coding sequences (CDS) without stop codon of *OsIAA3* (789 bp) and *OsARF16* (3165 bp) were amplified from wild-type and inserted into the binary vector pCAMBIA1301 driven by the ubiquitin promoter to generate *OsIAA3-OE* and *OsARF16-OE* plants, respectively. The genome editing of *OsIAA3* was performed using the CRISPR/Cas9 method as previously described (Ma et al., 2015). In briefly, the knockout target was screened by the web-based tool CRISPR-P 2.0 (http://crispr.hzau.edu.cn/CRISPR2/) to prevent potential off-target sites. Then, the single guide RNA (sgRNA) with target was introduced into pYLCRISPR/Cas9P_ubi_-H vector. The recombinant vector was introduced into *Agrobacterium tumefaciens* strain EHA105 to transform the Nipponbare cultivar. The mutants for *OsARF16* were purchased by BIOGLE (Changzhou, Jiangsu, China). The primers used in this study are listed in Supplementary table1.

### Protein sequence alignments

Protein sequence alignments were constructed using MAFFT version 7 (Katoh et al., 2019), and the aligned results were visualized using ESPript 3.0 (Robert and Gouet, 2014).

### NAA treatments

All the rice seedlings were grown in a greenhouse at 28℃ with 70% relative humidity and a 14 h light/10 h dark photoperiod at Wuhan University. To investigate the response of *OsIAA3* to auxin, 10-day wild-type seedlings were treated with Yoshida’s liquid culture solution containing 1 μM NAA, and the samples were collected at 0, 2, 4, 6, 8, 10 h after treatment. For the treatment of *osarf16* and wild-type, 10-day seedlings were treated with Yoshida’s liquid culture solution containing 1 μM NAA, and the seedlings were collected at 2 h after treatment.

### Protein degradation assays

Cell-free degradation assays were performed as described previously (Ma et al., 2023) with some modifications. The CDS without stop codon was fused to pET28a vector. The recombinant protein OsIAA3-His was expressed in the *Escherichia coli* Rosetta (DE3) strain and purified using the HisSep Ni-NTA MagBeads (Yeasen, Shanghai, China) according to the manufacturer’s protocol. Total protein from wild-type leaves was extracted using cell-free buffer (25 mM Tris-HCl, pH=7.5, 10 mM NaCl, 10 mM MgCl_2_, 5 mM DTT, 10 mM ATP, 4 mM PMSF). The mixture of OsIAA3-His and total protein was incubated at 4℃ for 1 h. After that, the mixture was treated with 0, 1, 10, 100 μM NAA at 30℃ for 3 h, and the reactions were stopped by adding loading buffer, and denatured at 98℃ for 10 min. The proteins were detected by anti-His and anti-Actin antibodies (ABclonal, Wuhan, China).

*In vivo* degradation assays were performed as described previously (Gao et al., 2021) with some modifications. The vector pCXUN-4×HA was used to generate OsIAA3-HA, and YFP-4×MYC (Hu et al., 2017) was used as a control. OsIAA3-HA and YFP-MYC were co-transformed into rice protoplasts. After incubation at 28℃ for 16-18 h in the dark, the total protein was extracted from the protoplasts using cell-free buffer (25 mM Tris-HCl, pH=7.5, 10 mM NaCl, 10 mM MgCl_2_, 5 mM DTT, 10 mM ATP, 4 mM PMSF). The total protein treated with 0, 1, 10, 100 μM NAA (with or without 100 μM MG132) were incubated at 30℃ for 3 h, followed by adding loading buffer, and denatured at 98℃ for 10 min. The proteins were detected by anti-HA and anti-MYC antibodies (MBL, Japan).

### RNA extraction, RT-PCR and qRT-PCR

Toal RNA was extracted from 1 cm young panicles, leaves or 10-day seedlings using Trizol reagent (Invitrogen, California, USA) according to the manufacturer’s instructions. Reverse transcription was performed using a Maxima H Minus First Strand cDNA Synthesis Kit with dsDNase (Thermo Fisher Scientific, Massachusetts, USA). qRT-PCR was performed by a Bio-Rad CFX96 Real-Time PCR system (California, USA) with Hieff^®^ qPCR SYBR Green Master Mix (Yeasen, Shanghai, China) according to the manufacturer’s instructions. Three independent RNA samples were used as biological replicates. The rice Os*ACTIN* gene (LOC_Os03g50885) was used as the internal control. The relative expression levels were calculated using the 2^−△△CT^ method.

### RNA-seq analysis

For RNA-seq analysis, total RNA was extracted from 1 cm young panicles of wild-type and *OsARF16-OE* transgenic plants with three biological replicates. Total RNA was sequenced by SeqHealth company (Wuhan, Hubei, China). The filtered clean reads were mapped to the rice Nipponbare reference genome (http://rice.uga.edu/index.shtml). Differentially expressed genes (DEGs) were identified with (|log2(FoldChange)| > 1 and pvalue<0.05) between wild-type and *OsARF16-OE* lines.

### Y2H assay

For the interaction between OsIAA3 and OsARFs in yeast, the CDS of *OsIAA3* and *OsARFs* were separately cloned into bait vector pGBKT7 and prey vector pGADT7 vector for Y2H assay. Yeast AH109 competent cells were co-transformed with OsIAA3 and OsARFs, and then plated on SD/-Trp/-Leu medium for 3 d at 30℃. The interactions were detected on high-stringency SD/-Trp/-Leu/-His/-Ade with or without X-α-Gal. The β-galactosidase activity was tested using a Yeast β-Galactosidase Assay Kit (Thermo Fisher Scientific, Massachusetts, USA) according to the manufacturer’s instructions.

For the homodimerization experiment, the full length and truncations of *OsIAA3* were cloned into both bait pGBKT7 and pray pGADT7 vector. The bait and prey plasmids were co-transformed into yeast AH109 strains. Transformants were cultured on SD/-Trp/-Leu and SD/-Trp/-Leu/-His/-Ade medium plate and incubated for 48-96 h at 30℃ for observation.

### Firefly LCI assay

The CDS of *OsIAA3* was cloned in-frame with the sequence encoding N-terminal half of firefly luciferase (nLUC) to generate nLUC-OsIAA3. The CDS of *OsARFs* were cloned in-frame with the sequence encoding C-terminal half of luciferase (cLUC) to generate cLUC-OsARFs. These recombinant plasmids were transformed into the GV3101 (pSoup-p19) chemically competent cells. Four different combinations shown in the figure were co-infiltrated on the same leaves of *Nicotiana benthamiana*. After 48-72 h, the leaves were treated with 1 mM Luciferin (Yeasen, Shanghai, China) reagent for 5-10 min in the dark, and images were captured using a chemiluminescence imager (Berthold, LB 985 NightSHADE).

### Subcellular localization and BiFC assay

The full length and truncations of *OsIAA3* were cloned into HBT-sGFP vector, driven by the CaMV 35S promoter. The empty vector was used as control. The BiFC was performed as described before with some modifications (Hu et al., 2012; Zhang et al., 2020). In briefly, the full-length CDS of *OsIAA3* was fused into pUC-SPYNE (the N-terminal end of YFP), and the full-length CDS of *OsARF16* was fused into pUC-SPYCE (the C-terminal end of YFP). Rice protoplasts were isolated from 10-day wild-type etiolated seedlings. Constructs were co-transformed into rice protoplasts in pairs. After incubation at 28℃ for 16-18 h in the dark, the high-resolution confocal laser-scanning microscope (TCS SP8, Leica) was used for observing GFP (488 nm), YFP (514 nm) and RFP (561 nm) fluorescence.

### Transcriptional activation analysis

To analyze the transcriptional activation of *OsARF16* in yeast, the full length and truncations of *OsARF16* were cloned into pGBKT7 vector. These vectors were co-transformed with pGADT7 into yeast strain AH109 competent cells in pairs.

For transcriptional activation in rice protoplasts, the CDS of *OsARF16* was cloned into GAL4DBD vector to generate GAL4DBD-OsARF16. The luciferase gene contained five copies of binding sites for GAL4 was used as a reporter, and the *Renilla luciferase* driven by the CaMV 35S promoter was used as an internal control. These plasmids were co-expressed in rice protoplasts (wild-type) and incubated at 28℃ for 16-18 h in the dark. The relative LUC activity (LUC/REN) was measured by a Dual-Luciferase^®^ Reporter Assay System (Promega, Wisconsin, USA) according to the manufacturer’s instructions.

### Cytological analysis

The paraffin section observation was performed as described previously (Liu et al., 2015b) with some modifications. The spikelet hulls were fixed before flowering with FAA (the ratio of 70% [v/v] ethanol, 40% [v/v] formaldehyde and glacial acetic acid was 18:1:1), and then dehydrated, embedded, sectioned, stained and observed.

The mature seeds were gold-coated using a sputter coater to facilitate the observation via scanning electron microscopy. The outer surface of the spikelet glumes was observed with a Hitachi S-3400N scanning electron microscope (Hitachi, Tokyo, Japan). Cell length and cell number were measured and calculated by ImageJ.

### Dual-luciferase transcriptional activity assay

The 2-kb promoter regions (before ATG) of *OsBUL1*, *DRW1*, *OsOPL1* and *OsSPL13* were cloned into pGreenII0800-LUC vector to generate reporters. The CDS of *OsARF16* was driven by the CaMV 35S to generate effector. Rice protoplasts (wild-type) were transfected with different combinations of plasmids and incubated at 28℃ overnight. The LUC/REN activity was measured by a Dual-Luciferase^®^ Reporter Assay System (Promega, Wisconsin, USA).

To detect the transcriptional activity in *Nicotiana benthamiana*, the recombinant plasmids were transformed into the GV3101 (pSoup-p19) and infiltrated into the leaves of *Nicotiana benthamiana*. After 48-72 h, the leaves were infiltrated with 1 mM Luciferin (Yeasen, Shanghai, China) reagent and incubated in the dark for 5-10 min before images captured by a chemiluminescence imager (Berthold, LB 985 NightSHADE).

### EMSA

The coding sequence of *OsARF16* DNA binding domain (114-237 aa) was cloned into the pGEX-6P-1 vector. The recombinant protein GST-OsARF16^114-237^ was expressed in the *Escherichia coli* Rosetta (DE3) strain and purified using the GSTSep Glutathione MagBeads (Yeasen, Shanghai, China) according to the manufacturer’s protocol. FAM-labeled and unlabeled primers were synthesized by GenScript (Nanjing, Jiangsu, China). For the DNA EMSA, the purified protein was incubated with probes in a binding buffer at 25℃ for 25 min, followed by separation on 5% native acrylamide gels in a 0.5× TBE buffer for 1.2 h. After electrophoresis, the gels were scanned by a Typhoon Trio Imager (GE, Amersham Typhoon).

## Supplementary Data

**Supplementary Figure S1**. Protein sequence alignment of OsIAA3 in Nipponbare and *osiaa3* knockout mutants.

**Supplementary Figure S2.** Subcellular localization and homodimerization of the truncated segments of OsIAA3.

**Supplementary Figure S3.** The interactions between OsIAA3 and partial OsARF repressors in yeast.

**Supplementary Figure S4.** Firefly LCI assay between OsIAA3 and partial OsARF repressors in *Nicotiana benthamiana*.

**Supplementary Figure S5.** BiFC analysis in rice protoplasts to detect the interaction between OsIAA3 and OsARF16.

**Supplementary Figure S6.** Protein sequence alignment of OsARF16 in Nipponbare and *osarf16* knockout mutant.

**Supplementary Figure S7.** Cytological analysis of glumes in Nipponbare, *osarf16* and *OsARF16-OE* plants.

**Supplementary Figure S8.** Relative transcriptional expression of several cell cycle-related genes in Nipponbare, *osarf16* and *OsARF16-OE* plants.

**Supplementary Figure S9.** Scatter diagram of DEGs between Nipponbare and *OsARF16-OE*.

**Supplementary Figure S10.** OsARF16 fails to directly regulate the expression level of *DRW1*, *OsOPL1* and *OsSPL13*.

**Supplementary Figure S11.** Relative expression level of *OsBUL1* in various organs and stages of Nipponbare.

**Supplementary Figure S12.** qRT-PCR analysis of *OsBUL1* in 10-day Nipponbare and *osarf16* seedlings with or without 1 μM NAA treatment.

**Supplementary Table1.** Primers and probes used in this study.

**Supplementary Table 2.** DEGs between wild-type and *OsARF16-OE* plants.

## Funding information

This work was supported by fund from the National Natural Science Foundation of China (31871592) and the Creative Research Groups of the Natural Science Foundation of Hubei Province (2020CFA009), and the Fundamental Research Funds for the Central Universities (2042022kf0015).

## Acknowledgements

Dr. Wei Huang from Guangzhou biomedshine Co.Ltd is highly appreciated for his kindly donation to this project.

## Author contributions

J.Hu. and F.X. designed this study. F.X., S.L., L.Z., Q.Z. and C.L. performed experiments. J.Huang. created the *osiaa3* mutants. B.X., Y.X. and X.Z. helped in identifying and harvesting the materials. J.Hu., F.X. and S.L. analyzed the data; F.X. and J.Hu. wrote the manuscript with feedback from all authors.

## Competing interests

None declared.

## Data availability

The data that support the findings of this study are available within the Figure and Supplementary Data and Tables, or are available from the corresponding author upon request.

